# Co-evolution of Yeast and Microalga: Identification of mutations that improve cooperativity

**DOI:** 10.1101/2024.06.04.597407

**Authors:** Jennifer R Oosthuizen, Rene K Naidoo-Blassoples, Debra Rossouw, Florian F Bauer

**Affiliations:** South African Grape and Wine Research Institute, Department of Agrisciences, Stellenbosch University, Stellenbosch 7600, South Africa

## Abstract

Laboratory-based evolution has long been successfully implemented for the generation of desired phenotypes in microbial strain development. The approach also provides insights into evolutionary mechanisms and adaptive molecular strategies which may be too complex to unravel in natural environments. The selection pressure in most of these approaches are physical or chemical factors or stressors, and only a few projects have attempted to use dynamic biotic selection pressures as a driver of evolution.

Here we investigate the formation of novel cooperative phenotypes between the yeast *Saccharomyces cerevisiae* and the microalga *Chlorella sorokiniana.* A synthetic ecology approach based on the cross-feeding of carbon and nitrogen was used to establish an obligate mutualism between these species which allowed for prolonged physical contact in a continuous co-culture system over 100 generations. Comparative genomic analysis of co-evolved yeast strains identified several potentially high impact Single Nucleotide Polymorphisms. Of these, two genes *ETP1* and *GAT1,* were found to synergistically contribute to the cooperative phenotype between yeast and microalgae These genes are involved in carbon (*ETP1*) and nitrogen catabolite (*GAT1*) repression with *ETP1* encoding a protein of unknown function, but implicated in ethanol tolerance and control of Hxt3p, while *GAT1* encodes a regulator of nitrogen catabolite repression. CRISPR generated null mutants of the parental (ancestral) yeast strain with either *ETP1*, *GAT1* or both genes deleted, were shown to mimic the co-evolved phenotype with improved cooperativity observed when paired with *Chlorella sorokiniana* suggesting a possible role of these genes in the establishment of mutualisms between yeast and microalgae.

**Importance:** Multispecies cultures have tremendous biotechnological potential but are difficult to control and show unpredictable population dynamics. This research aims to comprehensively characterise the behaviour and attributes of co-cultured microbial species, with the aim of optimising their combined functionality in a targeted manner. Taken together, our results demonstrate the importance and efficacy of thoughtfully integrating biotic selection pressures into strain development projects. The data also provide insights into specific molecular adaptations that favour cooperative behaviour between species. The co-evolutionary dynamics between *Saccharomyces cerevisiae* and other microbial species hold immense promise for unlocking novel insights into evolutionary biology, biotechnological applications, and our understanding of complex microbiological systems. Finally, the molecular characterisation of ecosystem-relevant traits provides significant impetus to the annotation of microbial genomes within an evolutionary relevant, multispecies context.

## Introduction

Laboratory evolution of microorganisms, such as yeast and microalgae, can provide a framework for the rapid adaptation of biotechnologically relevant species for specialised environments and biotechnological processes (Castledine et al. 2020). The strategy has primarily been applied to single species in pure cultures and using abiotic factors as selection pressures. Only a few projects have attempted to use biotic selection pressures, i.e., the presence of other species, to investigate adaptions within an ecological context, and large-scale phenotyping of strains has not been attempted in multispecies environments (Hom and Murray 2014; Naidoo et al. 2019; Oosthuizen et al. 2020; Venkataram et al. 2023). As a consequence, genome databases are almost devoid of ecosystem-relevant functional gene annotations. Moreover, the underlying mechanisms of such interactions over time are not well characterised, likely due to the difficulty of keeping species in prolonged physical contact for an extended period of time, without one species outcompeting the other. By using ecosystem engineering to create symbiotic states such as obligate mutualisms we can overcome this limitation (Venkataram et al. 2023b). Yeast and microalgae are not known to naturally form symbiotic relationships, although lichens may be considered as an evolutionarily relevant example of the potential of such associations. However, co-cultures of these microorganisms are potentially of significant ecological importance as they can shed light on how interspecific cooperative relationships are established (Lutzu and Dunford 2018; Oosthuizen et al. 2020). Synthetic symbiotic relationships between yeast and microalgae have been shown to involve intricate interactions and reciprocal benefit within synthetic ecosystems (Hom and Murray 2014b; Venkataram et al. 2023). Yeast can provide essential metabolites, such as vitamins and organic compounds, to support the growth and survival of microalgae, while microalgae offer a nutrient-rich environment and physical protection for yeast (Starmer and Lachance 2011). Understanding the co-evolutionary dynamics of such mutualistic associations can provide insight on the molecular mechanisms, genetic adaptations, and ecological implications involved in the long-term coexistence and co-evolution of the organisms involved (Mougi 2016). By delving deeper into co-evolutionary processes, researchers can unravel the evolutionary strategies, selective pressures, and potential trade-offs that shape the stability and functional diversity of yeast-microalgae mutualisms. This knowledge can contribute to broader insights into the coevolutionary dynamics of symbiotic interactions and their significance in various ecosystems. To explore these aspects, further studies employing genomic, physiological, and ecological approaches are needed. Although a significant body of work exists investigating both yeast and microalgae in monoculture settings, very little research exists on the two species applied in a symbiotic co-culture or its effects over time.

The research presented in this work using the model yeast *Saccharomyces cerevisiae* and an industrially relevant microalga, *Chlorella sorokiniana* show clear impacts of co-evolution over time on molecular targets which play a role in cooperative phenotypes. A continuous co-culture system and co-evolutionary platform, previously described by (Oosthuizen et al. 2020) has created a collection of co-evolved yeasts for further characterisation. Several yeast and microalgae co-evolved pairings with improved biomass production were selected for further study. Comparative genome analysis of the parental and several evolved *S. cerevisiae* strains revealed potentially high-impact single nucleotide polymorphisms (SNPs) in the evolved strains. Here we demonstrate that at least two of these SNPs, affecting the *ETP1* and the *GAT1* genes, individually and synergistically, contributed to the improved mutualism. This interplay between carbon and nitrogen metabolism might be key to the establishment of these mutualisms and their maintenance over time.

## Results

### Co-evolved strains show improved phenotypes in non-selective growth conditions

*Saccharomyces cerevisiae* and *Chlorella sorokiniana* were co-evolved for 100 generations and evolved strains of each species were paired and screened in an all-against-all manner (Oosthuizen et al. 2020). Co-evolved pairings were selected based on increased biomass accumulation of both species compared to the parental pairing when grown under selective growth conditions (obligate cross-feeding of carbon and nitrogen). Three pairings were retained for further investigation. Three co-evolved *S. cerevisiae* isolates, which had been co-evolved for 100 generations, referred to here as Y100.A, Y100.B and Y100.C were co-cultured with a co-evolved microalga and compared to the parental pairing. For these pairings, the co-evolved *C. sorokiniana* isolates all accumulated six times more cells than the parental strain over a period of four days (Fig.1A – C). All three yeast isolates therefore supported microalgal growth to the same extent. However, the yeast strains differed in their growth phenotypes. While all three reached significantly higher cell numbers over four days, final yeast cell concentrations differed significantly in the co-cultures (Fig.1D).

**Figure 1:**
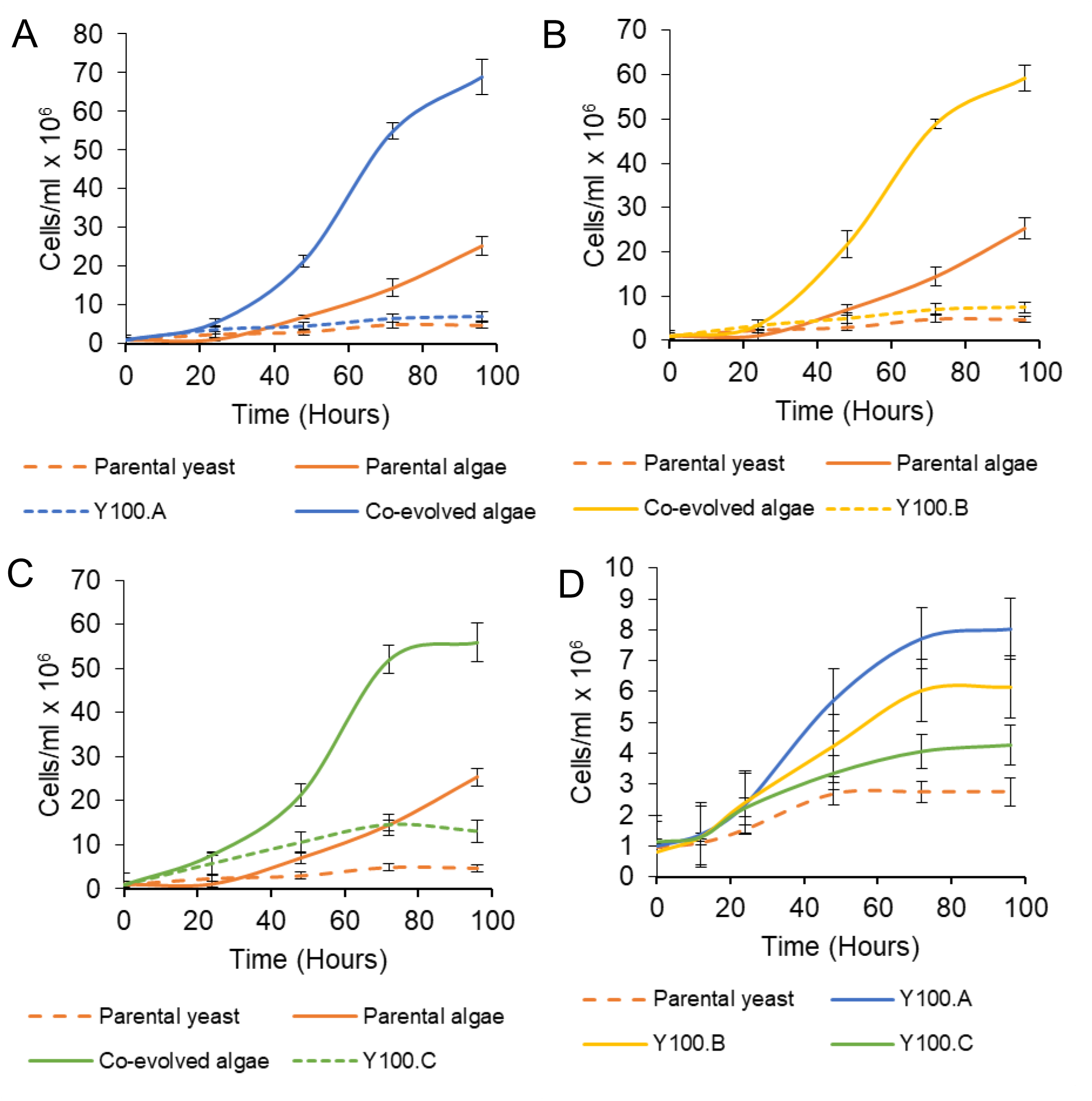
A-C represent co-culture growth of yeast and microalgae strains in cells/ml over time under selective conditions. *S. cerevisiae* growth is represented as a dotted line and *C. sorokiniana* growth is represented as a solid line. The parental pairing is represented in each graph (orange) along with Y100.A (blue), Y100.B (yellow) and Y100.C (green). Figure 1D shows the growth of the parental yeast (represented as a dotted line) and the three co-evolved *S. cerevisiae* strains in cells/ml over time under non-selective conditions in monoculture. Data represent the mean ± standard deviation (*n* = 3).

The co-evolved *S. cerevisiae* were compared to the parental strain in monoculture under non-selective conditions. To allow growth of the monocultures, nitrite was replaced by ammonia, as *S. cerevisiae* cannot utilise the nitrite provided under selective conditions. All co-evolved isolates grew to a higher cell count than the parental strain, with Y100.A reaching the highest cell concentration of 8,1 x10^6^ cells/ml (Fig.1D) compared to only 2,2 x10^6^ cells/ml of the parental yeast strain. Y100.C was chosen for further investigation based upon the improved co-culture phenotype (Fig. 1C). It was noted that although Y100.C accumulated the least biomass in monoculture under non-selective conditions (4 x 10^6^ cells/ml), it accumulated the most biomass in co-culture under selective conditions (6,7 x 10^7^ cells/ml) among the co-evolved isolates tested. These diverging phenotypes suggest that the three strains are genetically distinct.

### Identification of five potentially high impact SNPs present in co-evolved strains

The genomes of the co-evolved and parental *S. cerevisiae* strains were sequenced and assembled for variant analysis using *Saccharomyces cerevisiae* S288C (GenBank assembly accession: GCA_000146045.2) as the reference genome. Assembled genomes showed a 97% alignment to the lab strain reference genome of S288C and a sequencing depth of approximately 30x (Table 1).

**Table 1:** NGS reads and draft genome features for the parental and co-evolved *S. cerevisiae* isolates using S288C as the reference genome.

| Isolate | GC content (%) | Read alignment (%) | Draft genome |  |
| --- | --- | --- | --- | --- |
|  |  |  | Total length (Mbp) | Sequencing depth |
| S288C | 38.38 | N/A | 12.07 | N/A |
| Parental | 36 | 97.4% | 11.78 | 29.47 |
| Y100.A | 36 | 97.2% | 11.77 | 30.97 |
| Y100.B | 36 | 96.9% | 11.78 | 31.17 |
| Y100.C | 36 | 97.7% | 11.78 | 29.88 |

Variants were identified in the parental and three co-evolved strains using the S288C reference genome. Any variants present in the parental strain were filtered out of the co-evolved strain datasets as these had been present before co-evolution took place. Most mutation types identified in the isolates were synonymous mutations, followed by missense and frameshift mutations (Table 2). A breakdown on the type of identified SNPs in each co-evolved *S. cerevisiae* isolate can be found in the supplementary material (Fig. S1).

**Table 2:** Number of nonsynonymous SNPs identified in co-evolved strains after filtering out SNPs present in the parental strain.

| Co-evolved strain | Total nonsynonymous SNPs | Unique nonsynonymous SNPs | SNPs present in all sequenced co-evolved isolates |
| --- | --- | --- | --- |
| Y100.A | 120 | 8 | 47 |
| Y100.B | 138 | 11 |  |
| Y100.C | 134 | 14 |  |

Each strain contained a varying number of SNPs (Table 2). SNPs were filtered based upon depth (>20), quality score (>200) and only homozygous SNPs were considered. This resulted in 47 nonsynonymous variants called in 41 unique genes. Supplementary Table S1 contains all SNPs detected in all three sequenced co-evolved isolates that fall within these parameters. Of these variants, five had hypothesised high impact effects on gene function, with two of the SNPs impacting a single gene (Table 3) based on a SIFT analysis of the Variant Effect Predictor (Ng and Henikoff 2003). The SIFT analysis predicted that these SNPs corresponded to loss of function mutations, here referred to as a null mutation. The four genes with hypothesized high impact effects to gene function were *HNM1, PSP1, ETP1, GAT1.* Two variations are present on the *GAT1* gene which acts as a transcriptional activator of nitrogen catabolite repression genes. *HNM1* is a plasma membrane transporter for choline, ethanolamine, and carnitine, *PSP1* is an Asn and Gln rich protein of unknown function and *ETP1* is a protein of unknown function required for growth on ethanol. The presence of these SNPs was verified using Sanger sequencing and primer sets indicated in Table 5.

**Table 3:**
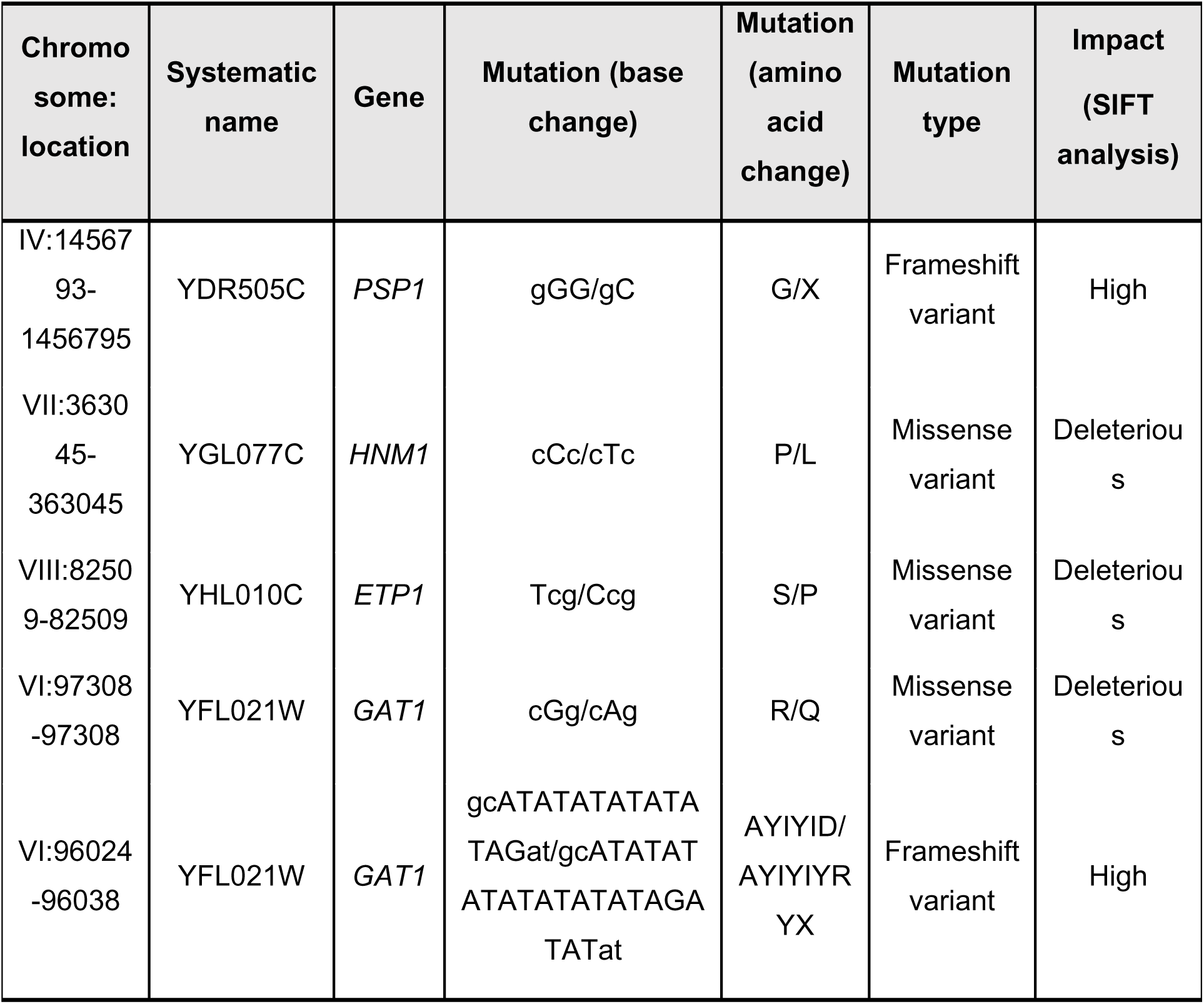
High impact or deleterious nonsynonymous variations identified in all co-evolved *S. cerevisiae* isolates sequenced. The location of the mutation, base change, amino acid change and mutation type are listed.

### Pairing co-evolved *C. sorokiniana* with Δ*etp*1 and Δ*gat1* deletion mutants result in improved biomass production

To evaluate the potential impact of these mutations, the Euroscarf BY4742 deletion library was used in a first screen to evaluate whether loss of function mutants of these genes impact co-cultures with *C. sorokiniana*. The parental and co-evolved *S. cerevisiae* (PY and Y100.C), BY4742 and the BY4742 deletion mutants were paired with the parental and co-evolved *C. sorokiniana* strain. Two of the pairings displayed growth identical to BY4742 paired with the parental and co-evolved *C. sorokiniana* (data not shown), while two of the mutant deletion strains (BY4742_Δgat1 S.C. and BY4742_Δetp1 S.C.) resulted in pairings with improved biomass production in line with the evolved phenotype (Fig.2).

**Figure 2:**
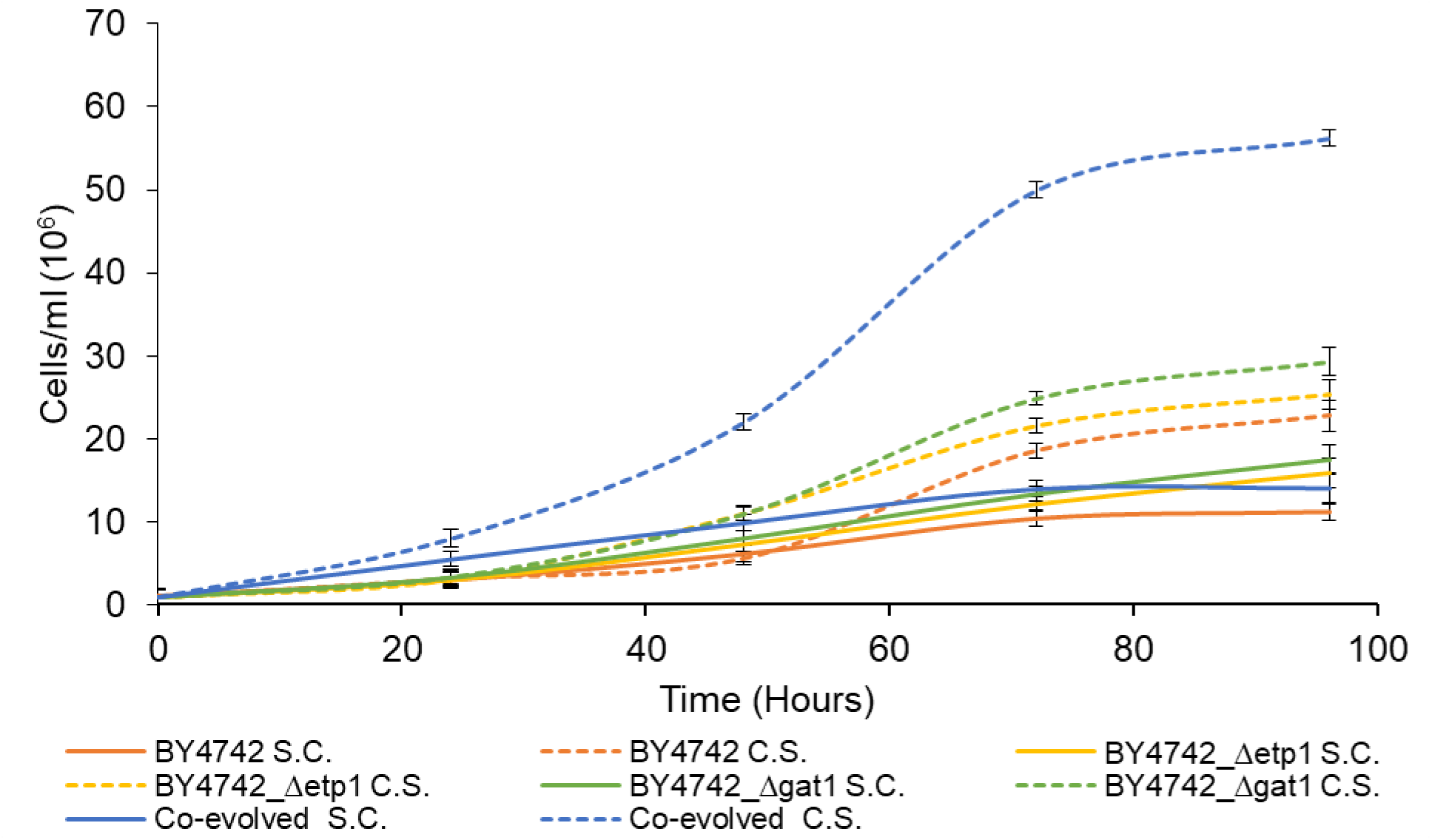
Growth of co-evolved *S. cerevisiae* Y100.C (blue), BY4742 (orange), BY4742_*Δetp1* (yellow) and BY4742_*Δgat1* (green) paired with co-evolved *C. sorokiniana* under selective conditions. S.C. denotes *S. cerevisiae* and is represented as a solid line and C.S. denotes *C. sorokiniana* and is represented as a dotted line. Data represent the mean ± standard deviation (*n* = 3)

### Inactivation of *ETP1* and *GAT1* in the parental strain results in improved co-cultures and show synergistic effects

The two genes of interest, *ETP1* and *GAT1* were inactivated in the parental strain (used for the co-evolution experiments) (PY*_Δetp1* and PY*_Δgat1*). In addition, a double deletion mutant, PY_*Δetp1*_*Δgat1*, was generated. The disrupted strains were generated using a CRISPR methodology as described by (Jakočiūnas et al. 2015), whereby a PAM sequence adjacent to the guide RNA recognition sequence on the antisense strand is replaced with nucleotides containing a stop codon (TAA) in the open reading frame (Fig. 3). The disrupted gene regions were confirmed by Sanger sequencing with primers provided in Table 6 and due to the diploid nature of the parental *S. cerevisiae* strain Sanger sequencing was also used to confirm a homozygous transformation as no double peaks were evident at the site of insertion (Fig. S2).

**Figure 3:**
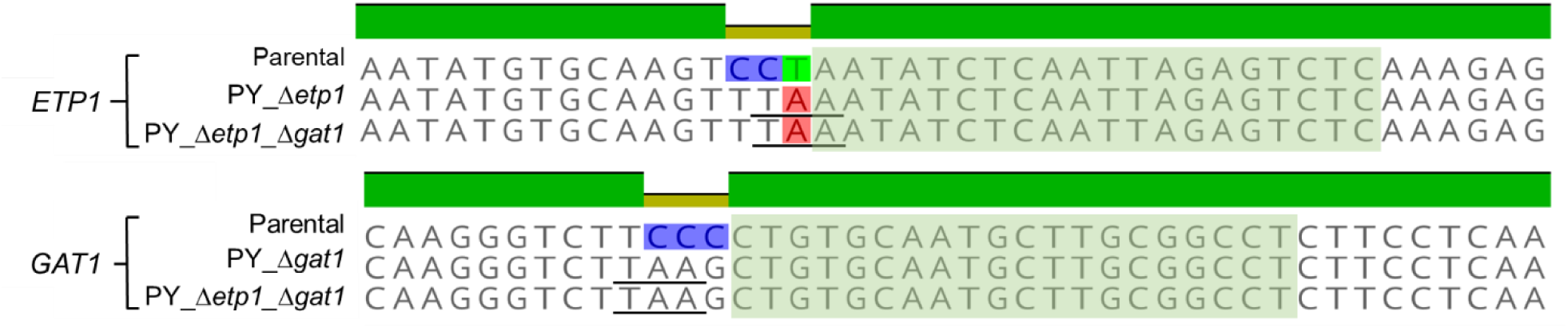
Alignment showing the introduction of a stop codon into the open reading frame of *S. cerevisiae* to replace the PAM sequence at the *ETP1* locus (top) and *GAT1* locus (bottom). The stop codon is underlined, and PAM sequences are highlighted in neon purple and green. gRNA recognition sequences are highlighted in light green. Aligned using Geneious Prime 2023.2.1.

The disrupted strains (PY_Δetp1, PY_Δgat1, and PY_Δetp1_Δgat1) were paired with both the parental and co-evolved *C. sorokiniana* strains to determine if their altered phenotypes differed from those seen with the parental pairing (Fig. 4A and 4B). The goal was to assess whether these genetic disruptions would lead to improved cooperative interactions and increased biomass accumulation compared to the parental pairing.

**Figure 4:**
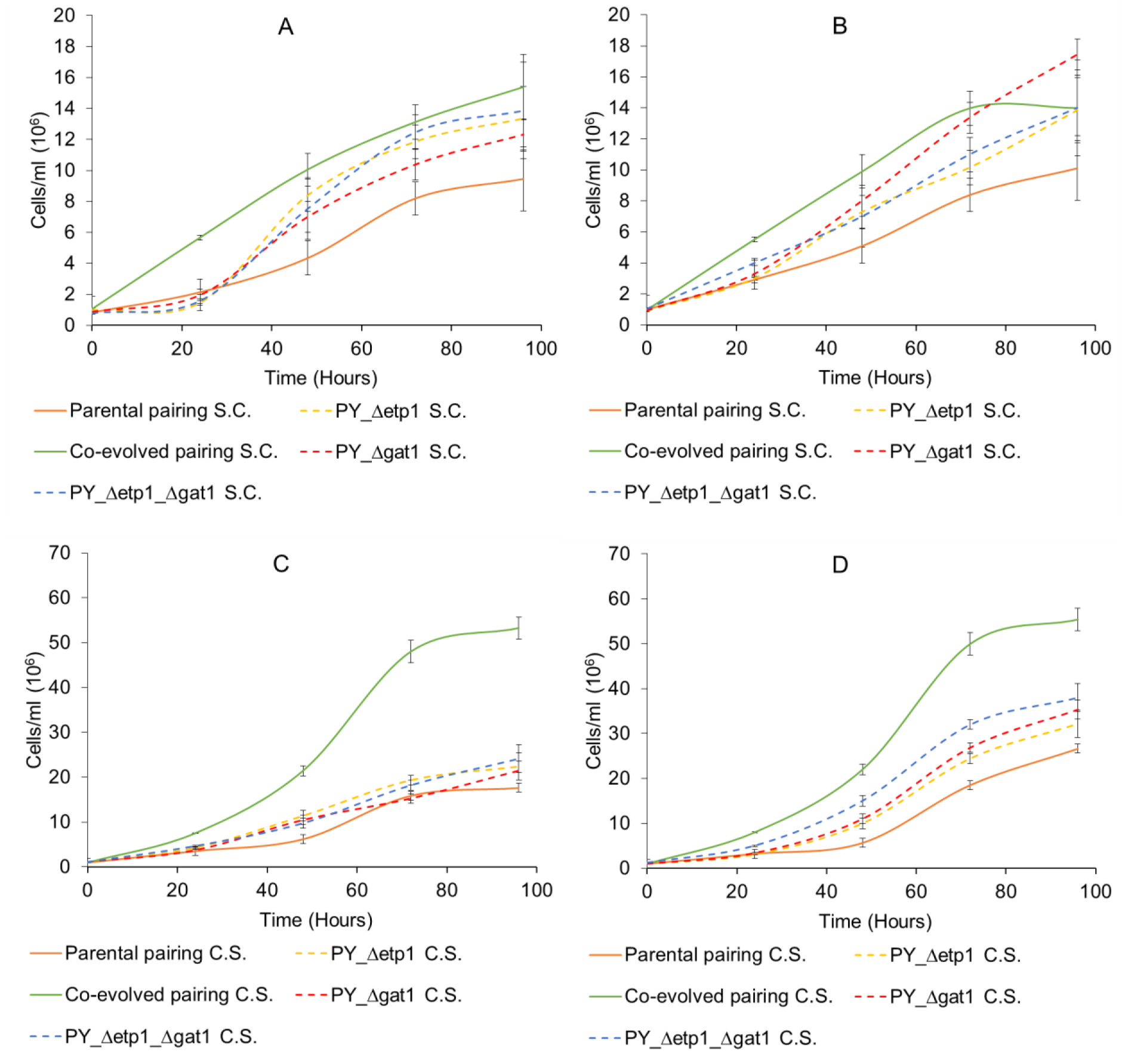
Growth of various *S. cerevisiae* strains in co-culture with *C. sorokiniana*. The parental *S. cerevisiae* is represented in orange, the co-evolved *S. cerevisiae* Y100.C in green, and the parental knock-out strains PY_Δetp1, PY_Δgat1, and PY_Δetp1_Δgat1 in yellow, red, and blue, respectively. Panel A displays yeast growth in co-culture with parental *C. sorokiniana*, while Panel B shows the yeast growth in co-culture with co-evolved *C. sorokiniana*. Microalgae co-culture growth is depicted in panel C (parental) and D (co-evolved) when paired with the same yeasts as in Panel A and B. In these graphs the parental and co-evolved strains are depicted with solid lines while the disrupted strains are represented as dotted lines. The data represent the mean ± standard deviation (*n* = 3).

The results clearly indicated that the disrupted yeast strains exhibited significantly enhanced cooperative growth when cultured with the co-evolved microalgae (Fig. 4B). Specifically, the single gene disruption strains (PY_Δetp1 and PY_Δgat1) showed more substantial biomass accumulation in the presence of co-evolved *C. sorokiniana* than with the parental strain. This suggests that even a single gene disruption can positively influence the cooperative dynamics between these species.

Moreover, the strain with both *etp1* and *gat1* genes disrupted (PY_Δetp1_Δgat1) demonstrated a particularly noteworthy phenotype. This strain exhibited growth and biomass accumulation characteristics that were much closer to those of the co-evolved phenotype than any of the other *S. cerevisiae* strains tested when in co-culture with the co-evolved microalgae. This indicates a synergistic effect when both genes are disrupted, leading to significantly enhanced cooperative growth (Fig. 4B and 4D).

Furthermore, the improved phenotype of the PY_Δetp1_Δgat1 strain was more pronounced when it was paired with the co-evolved microalgae (Fig. 4D) compared to the parental microalgae (Fig. 4C). This finding underscores the importance of co-evolution in enhancing the cooperative interactions between *S. cerevisiae* and *C. sorokiniana*. The co-evolution of both species appears to have led to mutual adaptations that optimize their growth and biomass production when grown together, suggesting that genetic modifications in both organisms contribute to their improved co-culture performance (Fig. 4B and 4D).

Overall, these results highlight the potential for using targeted genetic disruptions to enhance cooperative interactions and biomass production in mixed-species cultures, with significant implications for biotechnology and synthetic ecology.

### Inactivation of *ETP1* causes a change in the utilisation of non-preferred carbon sources

The parental, co-evolved, single and double mutated *etp1* and *gat1* strains were grown in media containing both preferred and non-preferred carbon sources in the form of glucose and mannose, respectively. The co-evolved and *etp1* mutant strains both grew to higher cell concentrations than the parental strain when provided with both glucose and mannose. The co-evolved strain reached the highest cell density at 7,1 x10^6^ cells/ml compared to 3,8 x10^6^ cells/ml of the parental strain (Fig.5A). Glucose utilisation in all four strains was similar in monoculture conditions reaching a final concentration of around 6 g/L after 36 hours of growth (Fig. 5B). Mannose utilisation however showed the parental strain only utilising around 2 g/L of mannose over 36 hours of growth with the co-evolved strain using 8 g/L over 36 hours of growth (Fig. 5C) and the *etp1* deletion strain using 6 g/L mannose. Mannose utilisation in the evolved and the mutated strains occurred throughout the period, including when glucose concentration was high at the beginning, suggesting an altered carbon utilisation pattern (Dynesen et al. 1998).

**Figure 5:**
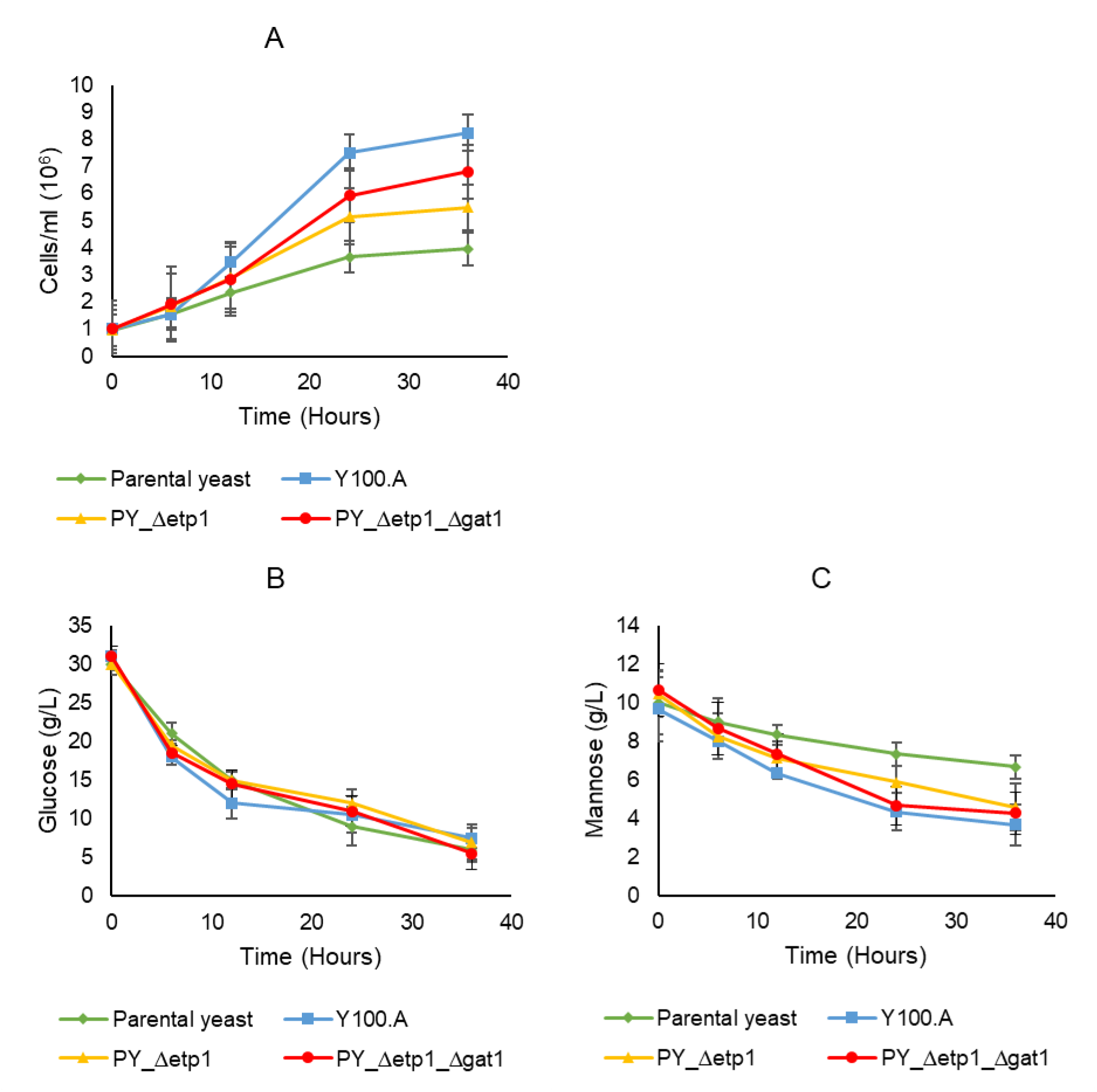
Graphs showing growth (A), glucose(B) and mannose (C) utilisation rates of the parental *S. cerevisiae* (green), co-evolved *S. cerevisiae* Y100.C (blue), PY_Δ*etp1* (yellow) and PY_Δ*etp1*_*gat1* (red) strains. Growth is represented in cells/ml (10^6^), Glucose concentration and mannose concentrations are represented in g/L. Data represent the mean ± standard deviation (*n* = 3).

### Inactivation of *GAT1* causes a change in the utilisation of non-preferred nitrogen sources

The parental, co-evolved, single and double disrupted *GAT1* strains were grown in media containing both preferred and non-preferred nitrogen sources in the form of ammonia and urea, respectively to evaluate whether nitrogen catabolite repression was affected by the co-evolutionary process. The co-evolved and Δ*gat*1 strains both grew to higher cell concentrations than the parental strain when provided with both glucose and mannose. The co-evolved strain however still reached the highest cell density at 6,2 x10^6^ cells/ml compared to 3,2 x10^6^ cells/ml of the parental strain (Fig. 6A). Ammonia utilisation in all four strains was similar in monoculture conditions reaching a final concentration of around 4 g/L residual ammonia after 36 hours of growth aside from the parental with 6 g/L remaining (Fig.6B). Urea utilisation rates however showed the parental strain only utilising approximately 3 g/L of urea over 36 hours of growth and the co-evolved strain using 7 g/L over the same period (Fig.6C) suggesting altered nitrogen source utilisation patterns compared to the parental strain. The continued use of the non-preferred nitrogen source indicates that a change in nitrogen catabolite repression may have taken place during co-evolution and that this phenotype is reproduced by *GAT1* disruption in the transformed strains.

**Figure 6:**
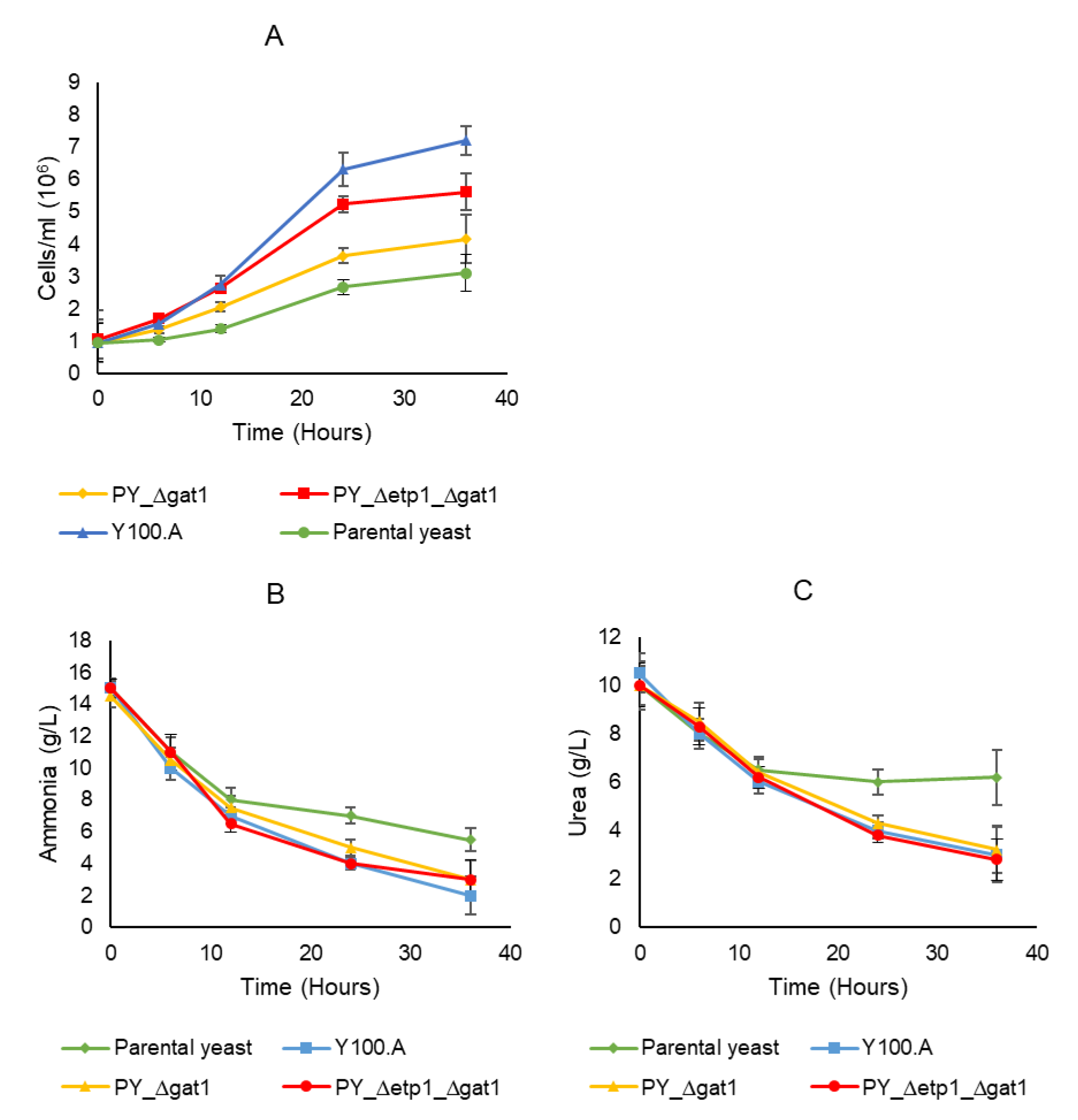
Graphs showing the growth (A), ammonia (B) and urea (C) utilisation rates of the parental *S. cerevisiae* (green), co-evolved *S. cerevisiae* Y100.C (blue), PY_Δ*gat*1 (yellow) and PY_Δ*etp1*_*gat1* (red) strains. Growth is represented as cells/ml (10^6^), and ammonian and urea in g/L. Data represent the mean ± standard deviation (*n* = 3).

## Discussion

In this study, an engineered mutualism between two species, yeast and microalgae, was used in a directed evolution set-up resulting in improved growth of both species in the co-culture. The co-evolutionary driver in this system is a synthetic obligatory mutualism based on the exchange of carbon and nitrogen sources between two species. On first sight, it may therefore not appear surprising that we identify two mutated genes in evolved *S. cerevisiae* strains that are involved in carbon catabolite and nitrogen catabolite repression as significant targets of the evolutionary adaptation. Indeed, the data show that these two mutations together account for a significant part of the co-evolved phenotype. However, the co-evolution takes place in an environment without any carbon or nitrogen sources that could trigger catabolite repression (mannose and nitrite) and the evolving yeast strains therefore are assumed to be completely de-repressed in these conditions. This suggests that the improved phenotype may not be directly linked to the role of these genes in catabolite repression, but to other functions of these genes.

The *ETP1* gene has been reported to contribute to increased ethanol tolerance of yeast, but our system is characterised by very low levels of ethanol, again making it unlikely that ethanol resistance is the trigger for the improved co-culture growth phenotype. Furthermore, existing data suggest that the overall contribution of *ETP1* to catabolite derepression and to ethanol resistance is minor. It is therefore likely that the phenotype is due to other functions of *ETP1*. Snowdon et al. 2009 showed that Etp1p is essential for the internalization and degradation of Hxt3p after switching from glucose to ethanol as the sole carbon source. It is plausible that disrupting the *ETP1* gene could lead to a slowed turnover of proteins, including the Hxt3p hexose transporters, perhaps providing an advantage for the uptake of mannose which is a non-repressing sugar (Dynesen et al. 1998). Guillaume et al. (2007) and Berthels et al. (2008) showed that a mutation in the *HXT3* gene enhanced fructose fermentation in *S. cerevisiae* and linked glycolytic flux to overexpression of *HXT3* while also suggesting a role in glucose and fructose sensing. Another possible explanation for the maintenance of this gene inactivation in co-evolved species is that the *ETP1* gene served no significant purpose under the conditions of continuous co-evolution, in which mannose is continuously provided, and that the loss of this gene did not negatively affect growth of the cell.

*GAT1*, on the other hand, is a bona fide activator of genes subjected to nitrogen catabolite repression (Hoffman and Winston 1987; Stanbrough et al. 1995; Hofman-Bang 1999). NCR enables yeast cells to preferentially utilize preferred nitrogen sources while repressing the utilization of less favorable nitrogen sources. Since the yeast is unable to utilise nitrite, all nitrogen required to support the increased biomass in co-cultures has to be supplied by the microalgae. Whether the microalgae naturally releases some nitrogen (amino acids, ammonium etc) during growth, or whether the yeast available nitrogen is released by dead microalgal cells, it will be available in very limited quantities, and whatever is released will likely immediately be taken up by the starved yeast cells. We were indeed unable to detect any nitrogen containing compounds (amino acids or ammonium) in the culture medium during co-culture (data not shown).

Our data show that the inactivation of *GAT1* in co-evolved *S. cerevisiae* impacts the NCR system and allows for the simultaneous utilisation of a wider range of nitrogen sources, enhancing rapid nitrogen uptake in nitrogen-limited environments. Hays et al. (2023) recently showed a similar phenomenon whereby fluctuating nitrogen starvation led to null mutations in the *GAT1* gene in 192 generations, approximately double the number of generations relative to this study (100 generations). Earlier studies have shown that proline, urea and allantoin utilisation in *GAT1* null mutants was increased compared to wild-type strains, and that *GAT1* null mutants display higher competitive fitness under poor nitrogen conditions (Hong et al. 2018). Taken together, our results likewise suggest that a reduction in the NCR leads to increased efficiency of uptake of non-preferred nitrogen in a low nitrogen environment. This adaptation enhanced competitive fitness in co-culture under selective conditions, enabling co-evolved *S. cerevisiae* to outcompete parental strains for limited nitrogen resources.

Our data therefore provide novel insights on gene function of a model species that has almost exclusively been studied in isolation or without consideration to the effects of interaction with other species over time. It also suggests that synergistic effects between *ETP1* and *GAT1* contribute to the co-evolved phenotype. Further investigation is needed not only for the characterization of *ETP1* and *GAT1* in *S. cerevisiae* monoculture but also in the context of co-culture. In *S. cerevisiae*, we observe some trade-offs, as demonstrated by the strain Y100.C exhibiting reduced growth in co-culture but improved growth in monoculture compared to other co-evolved strains and the parental isolate. The co-evolved strains have adapted in ways that improve ecosystem functioning under conditions in which these species were co-evolved, likely at the cost of optimal functioning in a wider range of environmental conditions. Gaining an understanding of how these processes impact species that are constantly engaging in interactions and undergoing evolution in their natural environment is crucial for comprehending the functioning and evolutionary dynamics of microbial species over time. The observed improvements in cooperative mechanisms such as carbon or nitrogen cross-feeding creates a more stable mutualistic relationship, in contrast to a scenario where perpetual growth occurs solely from positive feedback within the mutualism. Future work using carbon or nitrogen isotope tracing experiments could allow for the observation of carbon and nitrogen allocation within the system and provide insight as to how carbon and nitrogen metabolism might be rewired to improve co-operativity between yeast and microalgae. Though challenging, identifying relevant mutations in the co-evolved microalgae could provide insights into the microalgal side of the co-evolved phenotype which could help to understand the yeast phenotypes better.

The findings of this study shed light on the remarkable adaptability of microbial species in response to environmental challenges. Microorganisms, such as *Saccharomyces cerevisiae*, are adept at evolving novel genetic traits to thrive in diverse ecological niches. By disrupting genes crucial for carbon and nitrogen utilization, we witnessed the emergence of co-evolved strains displaying enhanced fitness under specific environmental conditions. This phenomenon underscores the inherent plasticity of microbial genomes and their capacity for rapid adaptation in dynamic environments. Understanding the mechanisms driving microbial adaptation holds immense significance, particularly in the realm of biotechnology and industrial applications. Microorganisms serve as invaluable resources for various biotechnological processes, including biofuel production, pharmaceuticals, and bioremediation. By deciphering the genetic underpinnings of microbial adaptation, we gain insights into optimizing microbial strains for enhanced productivity and efficiency in industrial settings.

The data presented here reveal molecular adaptations in response to biotic selection pressures which results in improved cooperativity between yeast and microalgae. Such ecosystem-relevant adaptations open new opportunities for the design of multispecies-based bioprocesses relying on stable associations of evolutionarily and phylogenetically unrelated partners. At the same time, these data open new perspectives on natural evolutionary processes. Indeed, biotic pressures are the main drivers of evolution in natural microbial ecosystems but have not been investigated extensively.

## Materials and methods

### Isolation and identification of microbial strains

Yeast, *S. cerevisiae*, and microalgae, *C. sorokiniana*, were isolated from winery wastewater sampled from a winery in the Stellenbosch wine region (Stellenbosch, South Africa) as described previously (Naidoo et al. 2019). Co-evolved strains were obtained from previous work in which *S. cerevisiae* and *C. sorokiniana* were continuously co-evolved for a period of 100 generations as described in (Oosthuizen et al. 2020).

*Saccharomyces cerevisiae* strain BY4742; BY4742Δ*gat1*; BY4742Δ*HNM1*; BY4742Δ*PSP1* and BY4742*Δetp1* (MATα; his3Δ1; leu2Δ0; lys2Δ0; ura3Δ0; thi4Δ0; EUROSCARF) were obtained from the culture collection of the South African Grape and Wine Research Institute (SAGWRI) at the University of Stellenbosch.

Freeze cultures (15% glycerol) were prepared for all strains grown in either YPD (*S. cerevisiae*) or TAP medium (*C. sorokiniana*) and were stored at −80 °C in order to pre-culture all strains from the same culture stock solutions for all experiments conducted.

### Preculture conditions

Yeast and microalgae pre-cultures were grown as monocultures in Tris–acetate phosphate (TAP) medium containing 1 × Hom’s vitamins (personal communication, Erik F.Y. Hom 2015) in 50 ml volumes until mid-log phase. Cell concentrations were quantified using a flow cytometer (CytoFLEX, Beckman-Coulter) for all experiments prior to inoculation at a cell density of 1 × 10^6^ cells/ml. All growth experiments were conducted in TAP media free of carbon and nitrogen sources at a temperature of 25 °C, agitation at 50 rpm with constant illumination unless stated otherwise.

### Continuous co-evolution and isolate screening conditions

Co-evolved isolates were obtained from co-evolution experiments performed by (Oosthuizen et al. 2020b). A continuous co-culture system was achieved by attaching a feed vessel to the bioreactor and modified TAP media containing 20 mM potassium nitrite (KNO_2_) and 2% mannose was fed into the bioreactor. These conditions resulted in an enforced mutualistic symbiosis between the species. Media was simultaneously pumped out of the bioreactor to maintain a constant volume of 600 ml in the bioreactor also using a peristaltic pump. The bioreactor remained clamped to create a closed system and was run for a total of 150 generations of co-culture.

### Isolate screening and growth conditions

Isolates were obtained from continuous co-culturing at approximately 100 generations as described in (Oosthuizen et al. 2020). These isolates were screened in an all-against-all manner under a variety of environmental conditions and three co-evolved *S. cerevisiae* isolates were identified for further investigation. Isolates were identified for further characterisation based on biomass production compared to the parental strain.

The obligate growth conditions used for co-culture growth of *C. sorokiniana* and *S. cerevisiae* was based on the cross-feeding of carbon and nitrogen. In this system, mannose is fermented by *S. cerevisiae* with the release of CO_2_ which is used as a carbon source by *C. sorokinian*a, and nitrite is metabolised to ammonium for use as nitrogen source by *S. cerevisiae*. As each microorganism is unable to grow on its own, semi-selective growth conditions were devised. Here, nitrite was replaced with ammonium to promote single culture growth of *S. cerevisiae*. The various C and N sources which were used to facilitate growth of mono- and co-cultures are listed below (Table 4).

**Table 4:** Monoculture and co-culture growth conditions for *S. cerevisiae* and C*. sorokiniana* used during this study.

| Species | Growth | Symbiosis | Media conditions | Carbon source | Nitrogen source |
| --- | --- | --- | --- | --- | --- |
| *S. cerevisiae* | Monoculture | N/A | Non-selective | 20 g/L mannose | 1.6 g/L ammonia |
| *S. cerevisiae*<br>(Nutrient utilisation testing - carbon) | Monoculture | N/A | Non-selective | 10 g/L mannose<br>30 g/L glucose | 1.6 g/L ammonia |
| *S. cerevisiae*<br>(Nutrient utilisation testing - nitrogen) | Monoculture | N/A | Non-selective | 20 g/L mannose | 15 g/L ammonia<br>10 g/L urea |
| *S. cerevisiae* and *C. sorokiniana* | Co-culture | Obligate | Selective | 20 g/L mannose | 1.7 g/L nitrite |

### DNA extraction

Genomic DNA was extracted from both the parental yeast strain and the co-evolved yeast isolate. The extraction was carried out from overnight cultures that had been cultivated at a temperature of 30°C in YPD broth. The isolation procedure involved mechanical cell disruption and subsequent phenol-chloroform extraction, following the method described by Hoffman and Winston in 1987. To concentrate the DNA, an ethanol precipitation step was employed. The resultant DNA was then resuspended in Milli-Q ultrapure water, and its quantity was determined using a NanoDrop® ND-1000 spectrophotometer from NanoDrop Technologies located in Wilmington, USA. To assess the purity of the DNA samples, a NanoDrop 8000 spectrophotometer manufactured by Thermo Fisher Scientific was utilized. This involved measuring the UV absorbance ratios at 260 to 280 nm (A260/280) and 260 to 230 nm (A260/230). Only DNA samples that exhibited purity within the ranges of 1.8 to 2.0 for the A260/280 ratio and 1.5 to 2.2 for the A260/230 ratio were considered suitable for further use. The quantification of DNA was conducted utilizing the Qubit dsDNA BR Assay Kit from Thermo Fisher Scientific, specifically using the Q32853 kit.

### Genome sequencing

The parental strain and the co-evolved isolate were sequenced at The Centre for Proteomic and Genomic Research (CPGR, Cape Town, South Africa).

Following the manufacturer’s instructions, the Nextera DNA Flex Library Prep Kit (Illumina, 20018704) was used to prepare the Illumina libraries. Using bead-linked transposomes, adapter sequences were added to fragmented DNA. It was then possible to separate the tagged DNA from the bead-linked transposomes. Individual libraries were supplemented with Illumina Nextera DNA CD Index using PCR (five cycles). Through the use of solid phase reversible immobilization, the indexed libraries were cleaned. The concentration of dsDNA in each library was measured using the fluorometric Qubit 1X dsDNA HS Assay Kit (Thermo Fisher Scientific, Q33231) after the individual indexed libraries had been purified.

According to the Nextera DNA Flex Library Prep Reference Guide, the 4 nM pooled sequencing library was denatured with 0.2 N NaOH, diluted to 12 pM, and mixed with the denatured PhiX positive control at a spike-in concentration of 1% v/v. The MiSeq Reagent Nano Kit v3 (600 cycles) (Illumina, MS-102-3003) was used to sequence the denatured library after it had been put onto the Illumina MiSeq instrument. The sequencer was set up to do a 2 X 301 cycle, paired-end, dual indexed sequencing run. At the conclusion of the run, FASTQ files were automatically created, saved to the MiSeq, and the Sequence Analysis Viewer (Illumina) program was used to evaluate the run’s quality.

### Genome assembly and variant analysis of NGS reads of co-evolved isolates

Sequencing data were uploaded to the Galaxy web platform and all data analysis was performed on the public server at usegalaxy.org (Afgan et al. 2018). Data analysis steps are summarized in Fig.7.

**Figure 7:**
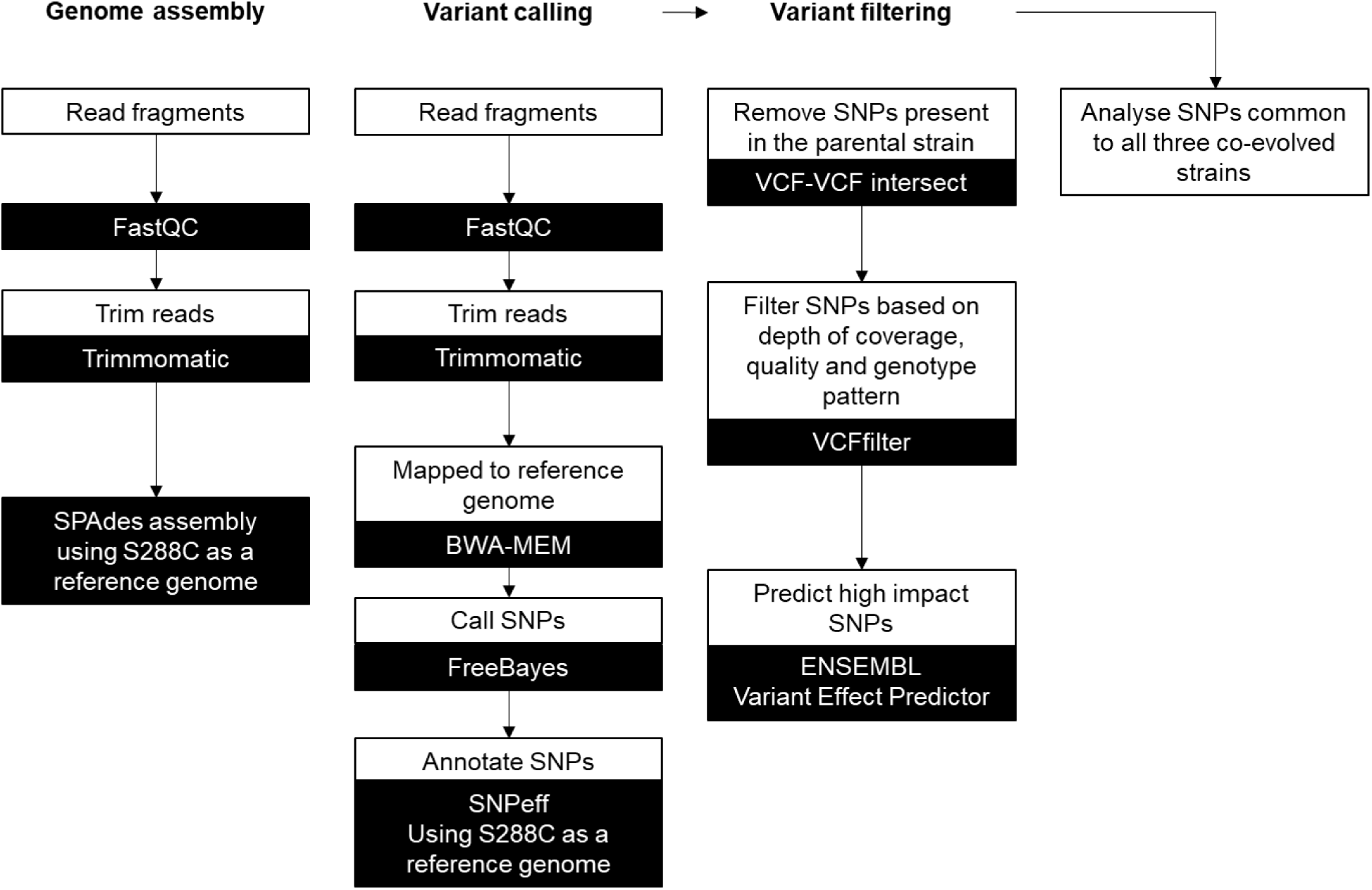
Pipeline for Next-Generation Sequencing (NGS) read analysis and variant calling is provided. The tools utilized within the Galaxy platform for this analysis are indicated in black. Specifically, BWA-MEM and snpEFF both employ the *S. cerevisiae* S288C genome (with GenBank assembly accession: GCA_000146045.2) as a reference.

To assess the quality of the sequence reads obtained from the CPGR, FastQC (version 0.72) developed by Andrews et al. (2010) was utilized. Based on the quality assessment, Trimmomatic (version 0.38.1) by (Bolger et al. 2014) was employed for various trimming tasks on the paired-end data. These operations included ILLUMINACLIP, which targeted the removal of Nextera adapter sequences from the reads, MINLEN to exclude reads shorter than 200 bp, and HEADCROP to eliminate the initial 20 bases from the start of the reads. For the purpose of mapping paired-end reads to the reference genome of *Saccharomyces cerevisiae* S288C (R64), BWA-MEM designed by (Li and Durbin 2010) was used. The trimmed paired-end reads were involved in the de novo genome assembly process, which was accomplished using SPAdes (version 1.0) as described by (Bankevich et al. 2012). To evaluate the quality of the genome assembly, Bandage Info (version 0.8.1) developed by (Wick et al. 2015) was employed.

BWA-MEM (Li and Durbin 2010)was used to map paired-end reads to the *S. cerevisiae* S288C (R64) reference genome (GenBank assembly accession: GCA_000146045.2). This was carried out for both the parental and co-evolved *S. cerevisiae* strains. Freebayes (Garrison and Marth 2012) was used to detect genetic variants in the strains compared to *S. cerevisiae* S288C. VCF-VCF intersect (Garrison et al., 2012) was used to identify single nucleotide polymorphisms found only in the co-evolved strains. These variants were filtered using VCFfilter (Garrison, 2012) based on a quality score greater than 500, a depth greater than 20 and only homozygous nonsynonymous SNPs were considered. The Ensembl Variant Effect Predictor (VEP) (Mclaren et al. 2016) was used to determine the effect of these variants on genes, transcripts and protein sequenced.

The sequences of the variant gene regions of *ETP1, PSP1, HNM1* and *GAT1* were confirmed by Sanger sequencing with primers listed in Table 5 on an ABI Prism 377 automated DNA sequencer at the Central Analytical Facility at Stellenbosch University.

**Table 5:**
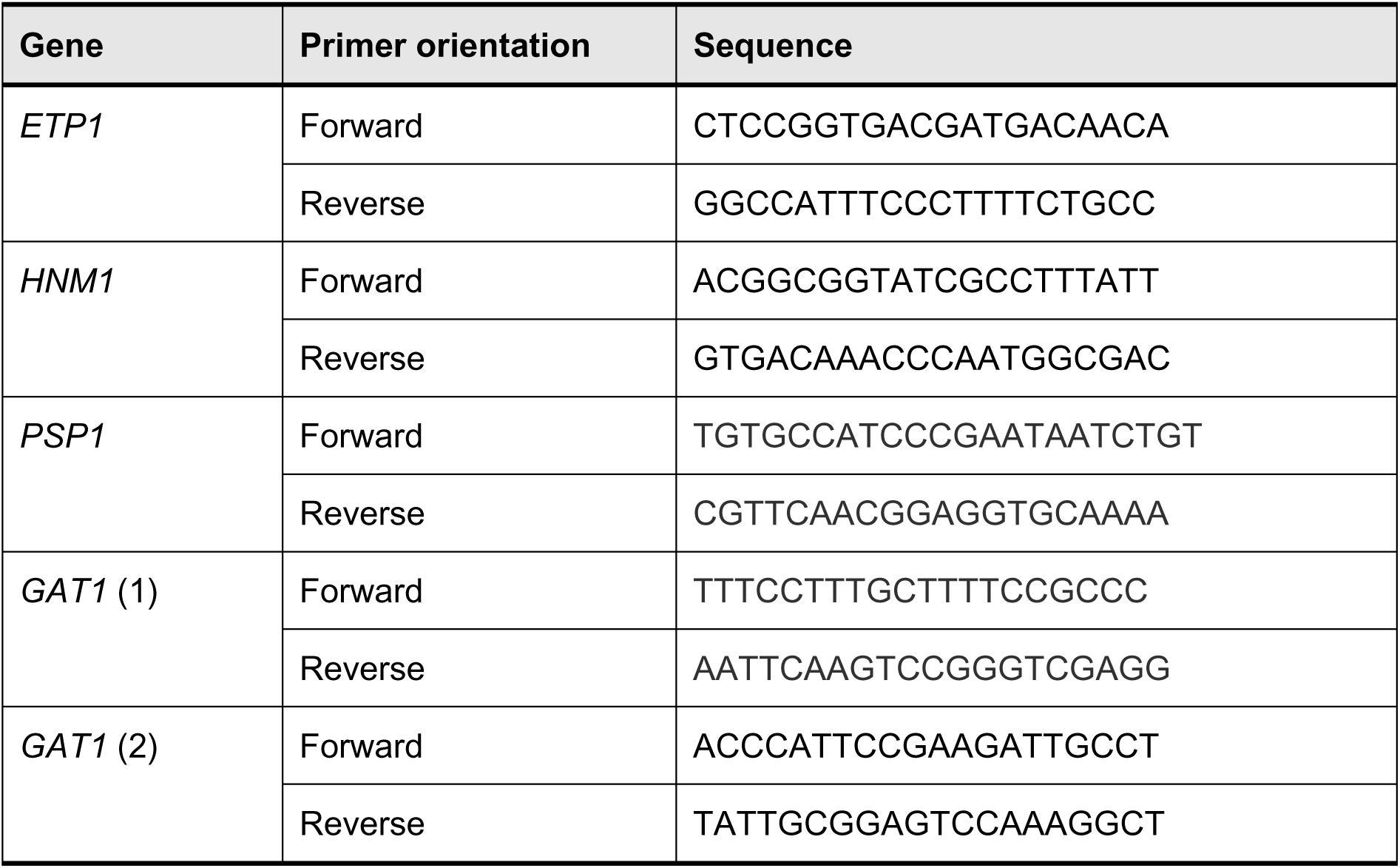
Primer sequences used in this study to confirm the presence of SNPs.

**Table 6:**
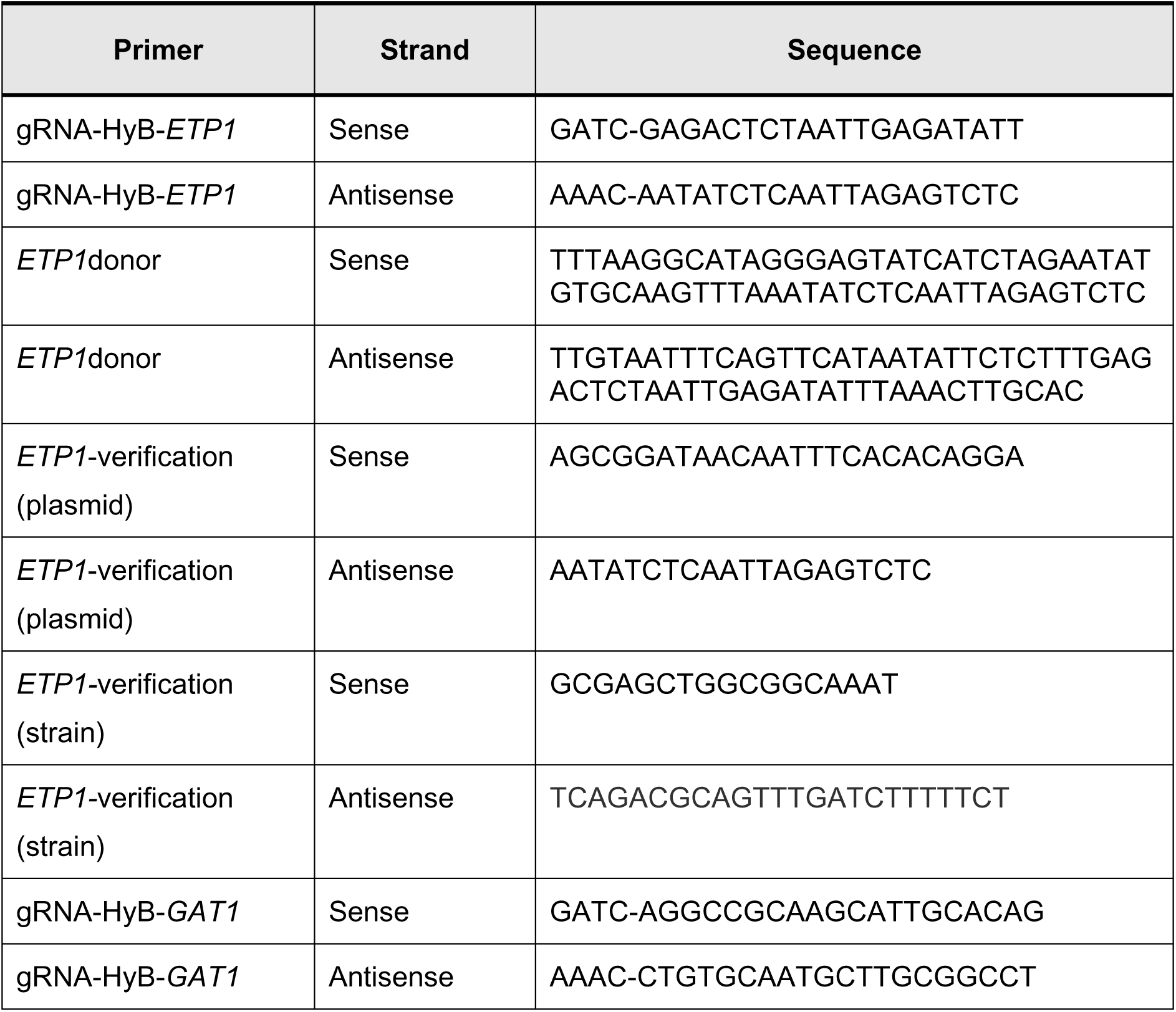
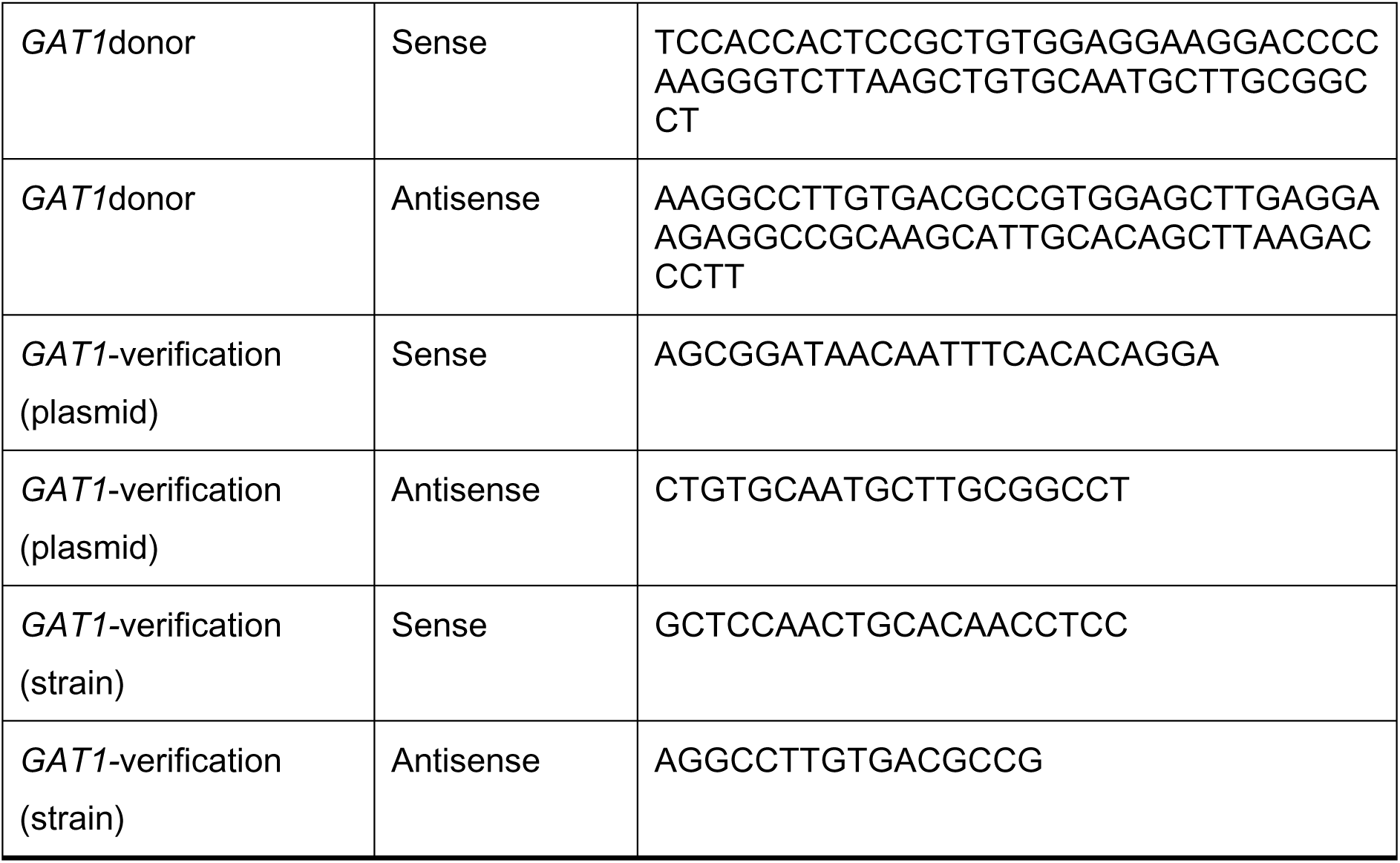
Primers used in this study for the generation of disrupted mutants of the parental yeast strain.

### Screening of deletion mutant pairings

Deletion mutant strains were chosen based on identified high impact SNP effects in co-evolved *S. cerevisiae* strain. Both the background strain and the deletion mutants were paired with the co-evolved *C. sorokiniana* strain for comparison to the parental and co-evolved pairings under selective conditions. Screening of co-cultures were conducted in 5 ml volumes of modified TAP media in 6-well cell culturing plates under continuous light (200 μmol m−2 s−1, SperAdvanced Light Meter 840022) with agitation (50 rpm). The plate was sealed using adhesive plate seals to create an airtight seal where CO_2_ can be trapped and, therefore, exchanged. Samples were taken every 24 h for a 96-h period. Growth, which was determined using CytoFLEX flow cytometer (Beckman-Coulter) using autofluorescence of *C. sorokiniana* cells to differentiate between species, was used as an indicator of altered phenotype.

### Yeast transformations to create null mutant

The creation of null mutants was carried out according to Zhang et al. (2014) whereby a stop codon is created by adding nucleotides to replace the PAM sequence.

#### Plasmid construction, propagation, and extraction

A custom gRNA-expressing plasmid was crafted following the gRNA synthesis protocol developed by Addgene Inc., with certain adjustments. The procedure encompassed two main steps. Initially, two distinct gRNA cassettes designed to target ETP1 and GAT1 were synthesized, with flanking sequences included for BSAI digestion. Subsequently, the synthesized gRNA DNA fragments were ligated into a plasmid named pRS42H after it had undergone BSAI digestion. This led to the generation of the plasmids gRNA-ETP1-HYB and gRNA-GAT1-HYB.

The maintenance and propagation of all plasmids (detailed in Table 7) involved the use of *Escherichia coli* DH5α from Thermo-Fisher Scientific, Waltham, MA, USA. These bacterial strains were cultured overnight at a temperature of 37°C on LB agar containing 100 μg/mL ampicillin. Following overnight incubation, bacterial colonies were introduced into LB media supplemented with 100 μg/mL ampicillin and incubated on a rotary wheel at 37°C overnight. This step was executed prior to extracting plasmid DNA.

**Table 7:** Plasmids used in this study for the generation of disrupted mutants of the parental yeast strain.

| Plasmid | Description | Marker | Source |
| --- | --- | --- | --- |
| pRS42H | Used to generate gRNA disruption cassette | Hygromycin B | Addgene Inc. |
| Cas9-NAT | Cas9 expression plasmid | Nourseothricin | Addgene Inc. |
| gRNA-*ETP1*-HYB | *ETP1* disruption gRNA cassette | Hygromycin B | This study |
| gRNA-*GAT1*-HYB | *GAT1* disruption gRNA cassette | Hygromycin B | This study |

For yeast strains, which were acquired from 40% (v/v) glycerol stocks stored at −80°C, cultivation was conducted on YPD agar (composed of 10 g/L yeast extract, 20 g/L glucose, 20 g/L peptone, and 20 g/L agar when needed). The cultivation medium was supplemented with 100 μg/mL nourseothricin and/or 200 μg/mL hygromycin B as required. Yeast cultivation was maintained at a temperature of 30°C for a period of 2 to 3 days.

The extraction of all plasmids was facilitated using the Purelink Quick Plasmid Miniprep kit from Thermo Fisher Scientific in accordance with the manufacturer’s guidelines. Both DNA and plasmids were purified employing the Monarch Genomic DNA Purification kit from New England Biolabs, Ipswich, Massachusetts, USA.

#### Yeast transformation

As the first step of transformation, plasmid Cas9-NAT (200ng per transformation) was transformed into the parental *S. cerevisiae* strain. Transformation was completed according to the method of Lin-Cereghino et al. (2005), with some modifications. Namely, competent cells were resuspended in 0.01 volumes of BEDS (10 mM bicine-NaOH, pH 8.3,3% (v.v^−1^) ethylene glycol, 5% (v.v^−1^) dimethyl sulfoxide (DMSO) and 1 M sorbitol) solution. The Gene Pulser® II electroporator (Bio-Rad Laboratories, Hercules, California, USA) was used: cuvette gap, 2.0 mm; charging voltage, 1500 V; resistance, 200 Ω; capacitance, 25 μF. After electroporation, 3 hours of recovery in yeast peptone dextrose sorbitol was sufficient and cells were collected by centrifugation at 1000 RPM for4 min, resuspended in 100 μl of saline (0.9% NaCl) and spread-plated. Transformed cells were plated onto selective medium containing 100 mg/L nourseothricin and grown for 3 days at 30°C until colonies formed.

The gRNA plasmid (∼500 ng) was then transformed together with the 90-mer donor dsDNA (∼2 μg) into the Cas9-expressing parental strain. Cells were plated onto medium containing 100 mg/L nourseothricin and 200 mg/L hygromycin B and allowed to grow for 2 to 3 days until transformants were ready to pick. The double-stranded 90-mer oligonucleotide repair DNA for either *GAT1* or *ETP1* disruption was generated using two rounds of PCR from two 60-mer primers (Table 6). These primers contain a region homologous to the upstream sequence of the PAM site and a 30-mer overlapping region encompassing the PAM sequence. A TAA stop codon was incorporated into the ORF to replace the PAM sequence to disrupt the relevant gene and simultaneously prevent repetitive Cas9 cleavage of the target site. To facilitate the removal of the gRNA plasmid strains were cultured in liquid YPD medium for a duration of 24 hours. Following this, the culture was streaked first onto a YPD plate to generate individual colonies. These colonies were subsequently transferred onto a YPD plate containing 100 mg/L nourseothricin via replica plating, thereby confirming the successful elimination of the plasmids. The integration of the stop codon was then confirmed by sequencing of the PCR product with primers found in Table 6 on an ABI Prism 377 automated DNA sequencer at the Central Analytical Facility at Stellenbosch University and sequences were analysed using Geneious Prime 2023.2.1.

### Nutrient utilization analysis of transformed strains

Carbon utilisation rates were evaluated using TAP media supplemented with both preferred carbon sources (glucose) and non-preferred carbon sources (mannose) at a concentration of 30 and 10 g/L respectively. Mannose and glucose consumption was monitored using the d-mannose-d-fructose-glucose Assay Kit from Megazyme (Bray, Ireland). These tests were conducted to establish how strains utilise carbon sources before and after co-evolution. Nitrogen utilisation rates were tested using TAP media supplemented with both preferred carbon sources (ammonia) and non-preferred carbon sources (urea) at a concentration of 15 and 10 g/L respectively. Nitrite consumption was monitored using the Griess Reagent Kit for nitrite quantification from ThermoFisher Scientific (Waltham, Massachusetts). Ammonia and urea concentrations were measured using Megazyme Urea/Ammonia Assay Kit (Bray, Ireland). These tests were carried out to establish how strains utilise nitrogen before and after co-evolution. Carbon and nitrogen concentrations were analysed in samples taken every 24 h. Volumes of 2 ml were centrifuged at 20 000 g for 3 min. The supernatant was then removed and stored at -20 °C until the assay was performed according to the protocols provided.

## Data availability

All supporting sequencing read data generated have been submitted to the National Centre for Biotechnology Information (NCBI) Sequence Read Archive (SRA) under BioProject no. PRJNA998225.

## Acknowledgements and contributions

F.F.B. and R.K.N.-B. conceptualized the study. J.R.O. designed and executed all experimental work and bioinformatic analyses. J.R.O., D.R., R.K.N.-B., and F.F.B. interpreted the data. J.R.O. wrote the manuscript. D.R., R.K.N.-B., and F.F.B. edited the manuscript. All authors approved the final manuscript.

This work was supported by the National Research Foundation (NRF) of South Africa through SARChI grant UID 83471 to F.F.B.

## Supplementary

**Table S1:**
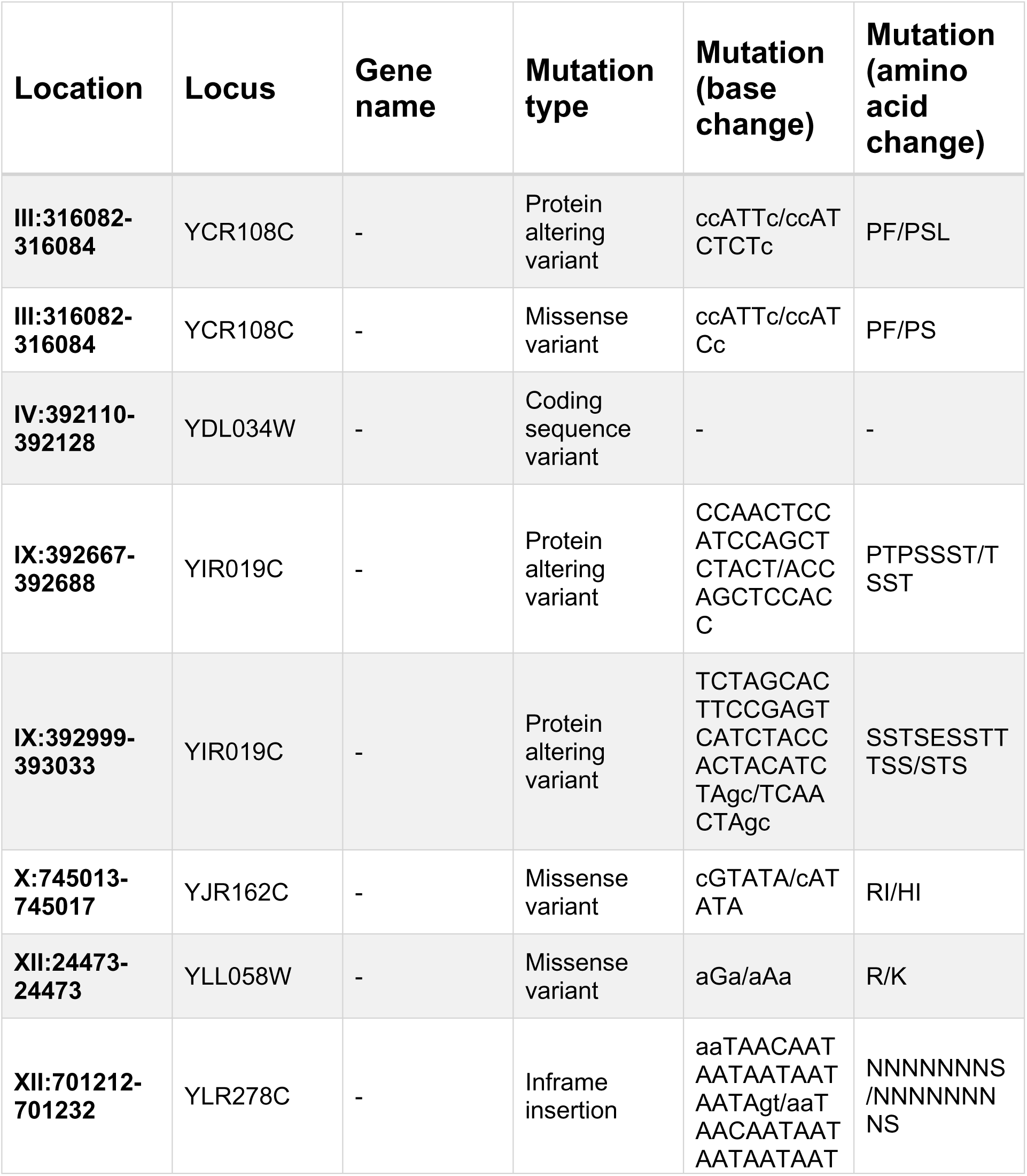
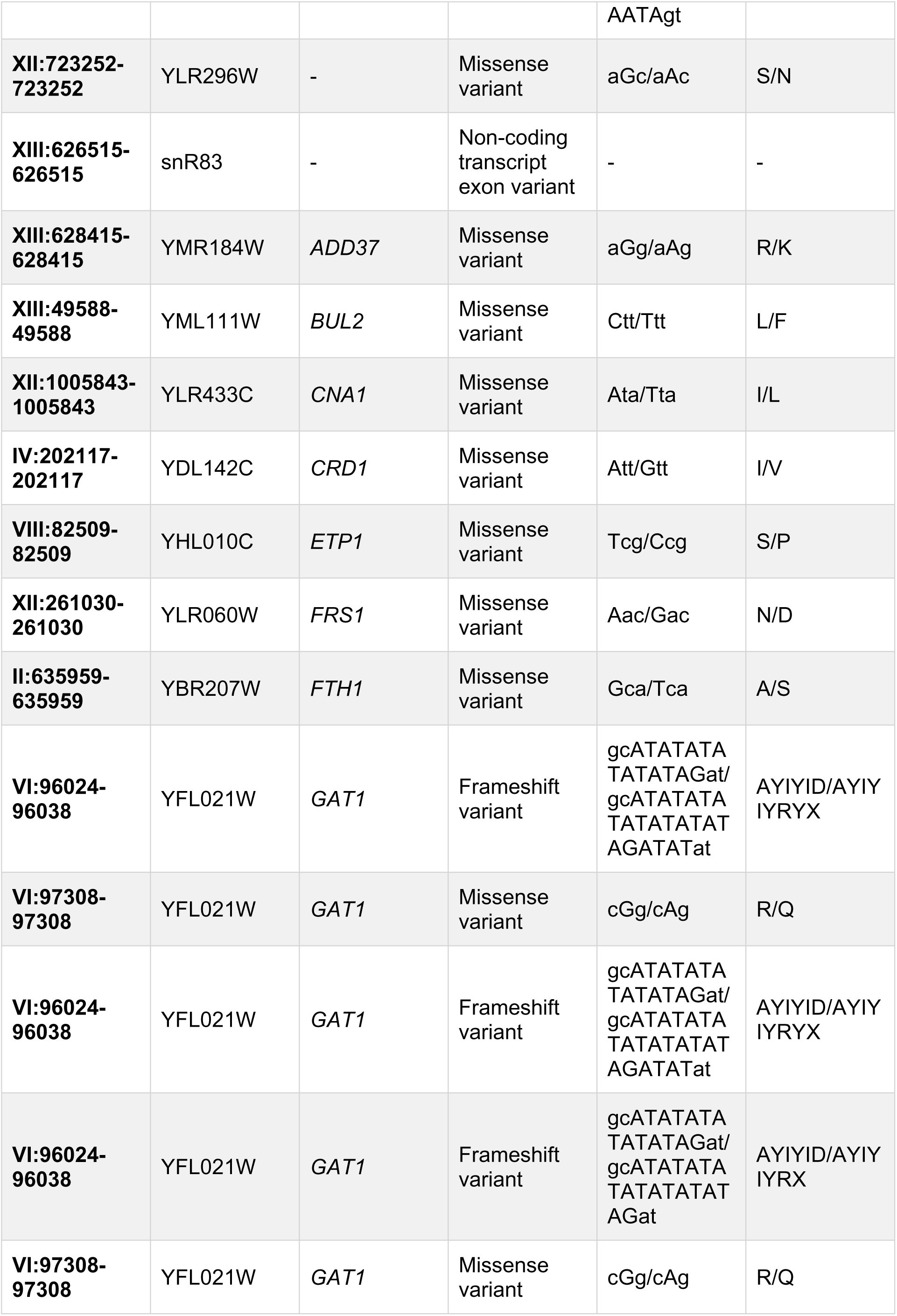
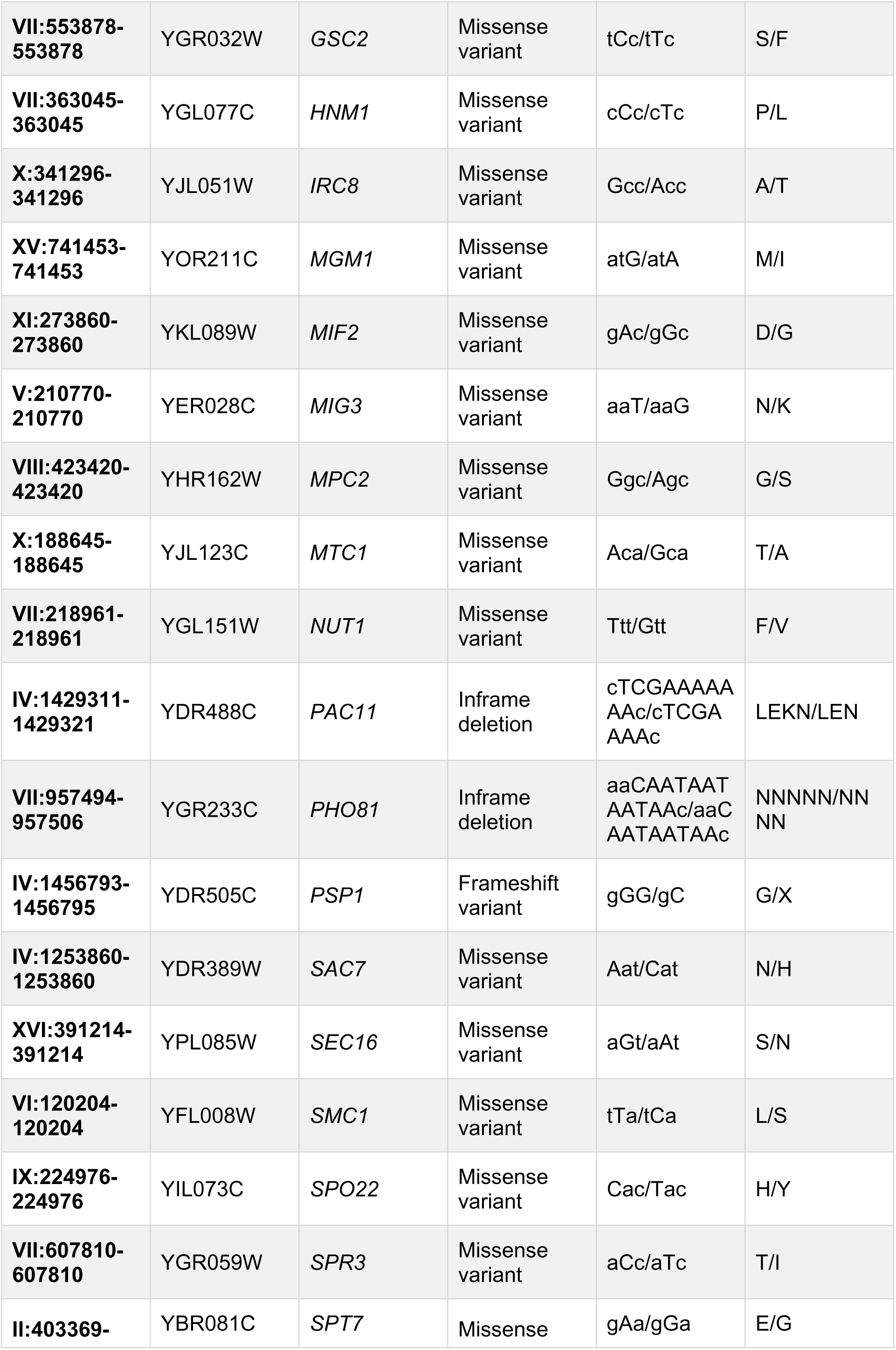
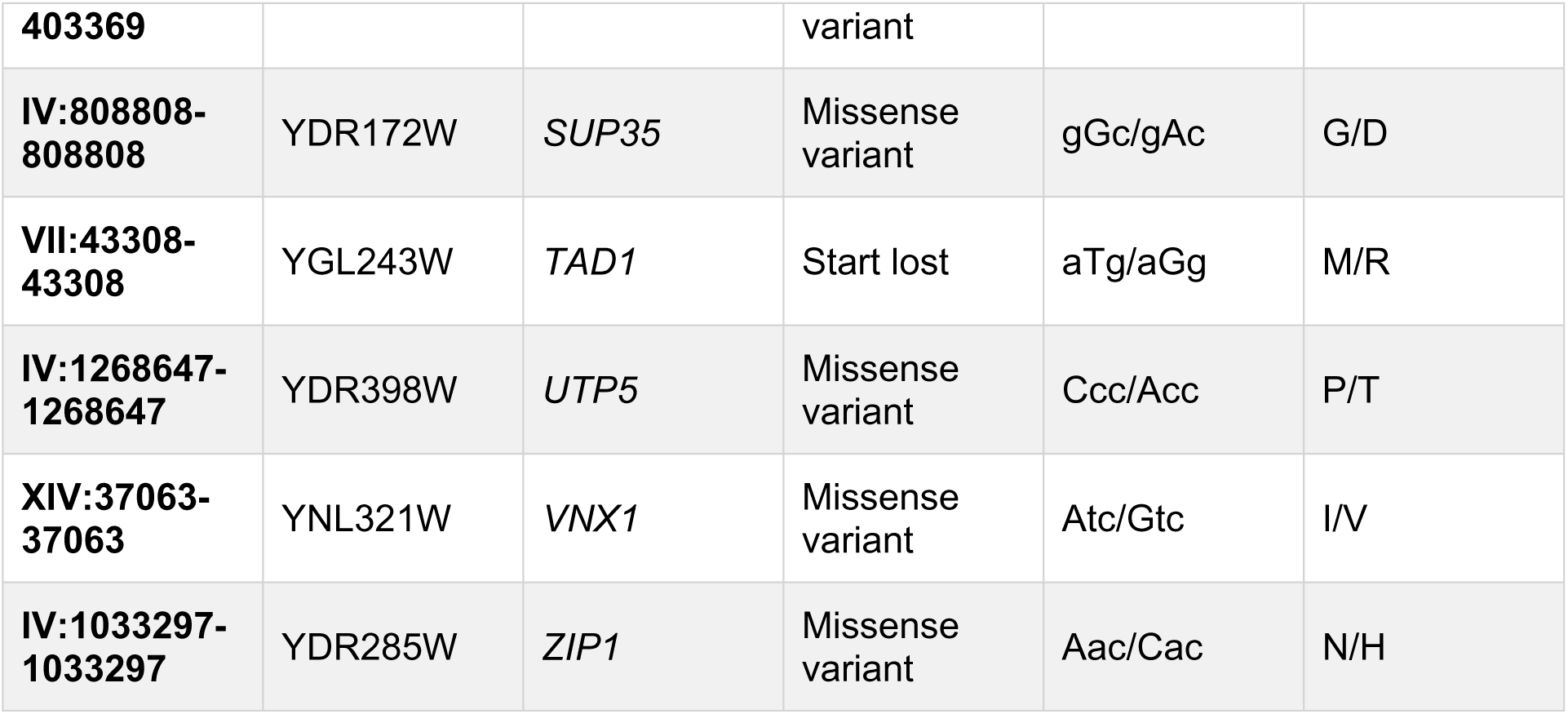
Nonsynonymous variations identified in all three co-evolved *S. cerevisiae* isolates. The location of the mutation, base change, amino acid change and mutation type are listed.

**Figure S1:**
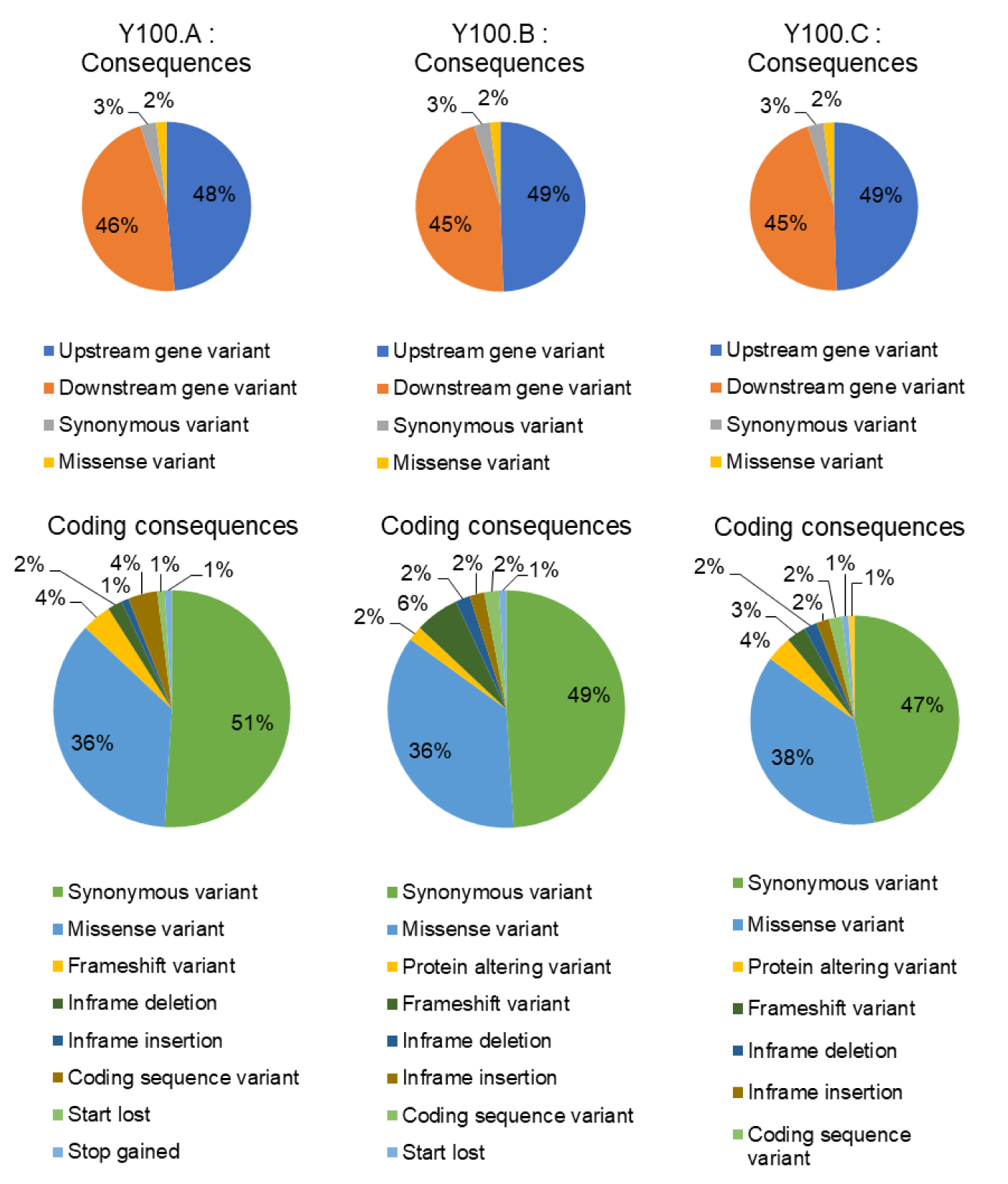
Variant Effect Predictor results indicating all consequences (top row) and specifically coding consequences (bottom row) of co-evolved strain variants in Y100.A, Y100.B and Y100.C.

**Figure S2:**
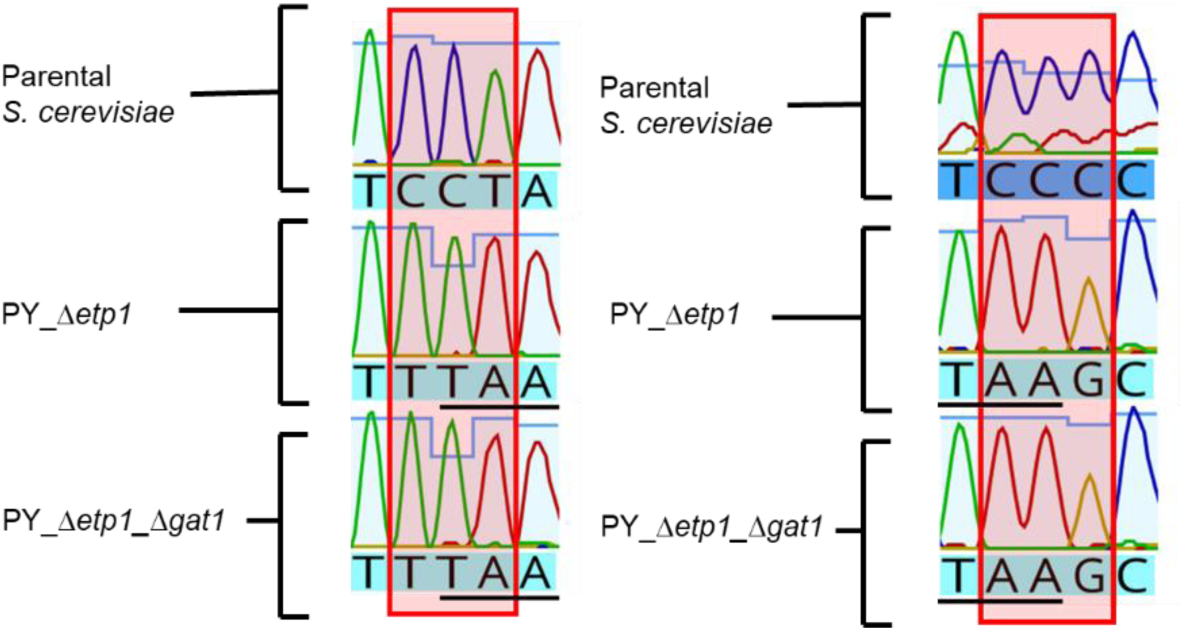
Sanger sequencing of region containing the inserted stop codon. Inserted codons are highlighted in red and stop codons in the open reading frame are underlined.

